# Island species-area relationships in the Andaman islands emerge because rarer species are disproportionately favored on larger islands

**DOI:** 10.1101/857672

**Authors:** Leana D. Gooriah, Priya Davidar, Jonathan M. Chase

## Abstract

The Island Species-Area relationship (ISAR) describes how the number of species increases with increasing size of an island (or island-like habitat), and is of fundamental importance in island biogeography and conservation. Here, we use a framework based on individual-based rarefactions to infer whether ISARs result from random sampling, or whether some process are acting beyond sampling (e.g., disproportionate effects and/or habitat heterogeneity). Using data on total and relative abundances of four taxa (birds, butterflies, amphibians and reptiles) across the Andamans and Nicobar archipelago, we examine how different metrics of biodiversity (total species richness, rarefied species richness, and abundance-weighted effective numbers of species emphasizing common species) vary with island area. Total species richness increased for all taxa, as did rarefied species richness for a given sampling effort. This indicates that the ISAR did not result because of random sampling, but that instead, species were disproportionately favored on larger islands. This disproportionate effect was primarily due to changes in the abundance of rarer species, because there was no effect on the abundance-weighted diversity measure for all taxa except butterflies. Furthermore, for the two taxa for which we had plot-level data (lizards and frogs), within-island *β* -diversity did not increase with island size, suggesting that heterogeneity effects were unlikely to be driving these ISARS. Overall, our results indicate that the ISAR of these taxa is most likely because rarer species are more likely to survive and persist beyond that which would have been expected by random sampling alone, and emphasizes the role of these larger islands in the preservation and conservation of species.

## Introduction

The Island Species-Area relationship (ISAR) describes the relationship between the number of species on an island and the area of that island, and has served as a basis for some of the most important theories in biodiversity studies, such as the theory of island biogeography (MacArthur and Wilson 1967, Warren et al. 2015). While the general pattern and shape of the ISAR is generally positive and its shape is described by a few key parameters (e.g., Triantis et al. 2012, Matthews et al. 2016), there remains uncertainty about the mechanisms underlying the ISAR and how they shape it (e.g., Chase et al. 2019). A deeper understanding of these mechanisms will not only provide insight into the processes that shape biodiversity and its variation on islands, but will also be important for devising plans for conserving biodiversity on islands, which house a disproportionate amount of diversity compared to their land area, but are also disproportionately influenced by human impacts and global change (Vitousek et al. 1997, Tershy et al. 2015).

The simplest explanation leading to the positive ISAR is random sampling—where larger islands have more individuals and as a result, a higher likelihood of passively sampling more species from the regional pool than smaller islands (Connor and McCoy 1979). Coleman (1981) provided an analytical approach to evaluate this null model, which Coleman et al. (1982) subsequently tested with bird abundances on islands, finding that they did not reject the random sampling hypothesis. Indeed, when appropriate data were available, random sampling has been implicated in a number of empirical studies of ISAR patterns (e.g., Haila 1983, Hill et al. 1994, Ouin et al. 2006, Bidwell et al. 2014, Gooriah and Chase in revision), though other studies have rejected the random sampling hypothesis (e.g., Ranta and As 1982, Bolger et al. 1991, Schoereder et al. 2004, Wang et al. 2010, Xu et al. 2017).

If the random sampling effect is rejected, two classes of biological mechanisms beyond random sampling can be invoked. First, island size can disproportionately influence some species relative to others (when random sampling is operating, effects are proportional); Connor and McCoy (1979) called these ‘*area per se’* effects to indicate that island area itself influences the relative abundances and likelihood of co-occurrence among species, and Chase et al. (2019) more generally called these ‘disproportionate’ effects. One prominent mechanism leading to disproportionate effects is the colonization-extinction dynamics inherent to MacArthur and Wilson’s (1963, 1967) theory of island biogeography. Likewise, population-level processes (Allee-effects or demographic stochasticity), which tend to be more pronounced on smaller rather than larger islands, can also lead to disproportionate effects.

Second, an increasing number of habitats, or an increase in habitat heterogeneity, with island area can also lead to more species on bigger islands (Kohn and Walsh 1994), particularly if species require specific or multiple habitat types (Williams, 1964, Hart and Horwitz, 1991, Guadagnin and Malchik 2007). However, disentangling disproportionate effects from habitat diversity can prove to be quite challenging as they can easily be confounded (Connor and McCoy 1979, Gilbert 1980, Boecklen and Gotelli 1984, Kohn and Walsh 1994); that is, bigger islands tend to have more diverse habitats (Hortal et al 2009). Furthermore, it is possible that area and habitat diversity together can better explain the variation of species patterns across islands (Ricklefs and Lovette 1999, Davidar et al. 2001, Triantis et al. 2003, Kadmon and Allouche 2007). Even within the same island archipelago, it is possible that different mechanisms underlie the response of different taxa to island area, depending, for example, on their dispersal capacity. For example, in a study of the ISAR of Caribbean islands, Ricklefs and Lovette (1999) suggested that birds were more likely responding to area alone, while habitat diversity effects were stronger for butterflies, amphibians and reptiles.

In this study, we use previously collected abundance data from four taxa that differ in their dispersal capacity—birds, butterflies, frogs and lizards—from the Andaman and Nicobar archipelago in the Bay of Bengal to examine the possible mechanisms underlying their respective ISARs. For birds and butterflies, we were able to explicitly test the null hypothesis of random sampling against more ecological mechanisms underlying the ISAR of these taxa using the individual-based rarefaction framework outlined in Chase et al. (2019). For frogs and lizards, we additionally had spatially-explicit plot level data, which allowed us to additionally test the potential role of habitat heterogeneity.

## Material & Methods

### Study site and sampling methods

The Andaman and Nicobar archipelago includes 556 islands, islets and rocks and is made up of four large contiguous regions: North, Middle, Baratang and South Andamans forming of over 5000 km^2^ in total area, surrounded by many isolated islands. The forest types across islands are diverse, ranging from evergreen forests to deciduous forests and mangroves (Champion and Seth, 1968, Davidar et al 2002). Bird and butterfly surveys were carried out on 38 and 25 of these islands respectively (varying in size from 0.03 to 1375 km^2^) in 1992 as part of the studies by Davidar et al. (1996) and Devy et al. (1998); data on the abundances of species from these surveys were previously unpublished (provided here in the Appendix). Frog and lizard surveys were carried out on 15 of these islands (varying in size from 0.03 to 1375 km^2^) between 2010-2012 and were previously published by Surendran and Vasudevan (2015).

Transect methods were used to sample forest birds and butterflies (for more details, see Davidar et al. 1996, Devy et al. 1998). Bird sampling was conducted between 1992-1994 during the dry seasons, along 1 km length transects laid within each habitat type on the bigger islands. On smaller islands, transects cut through all the habitat types. The number of transects placed increased with the size of that habitat. Butterflies were sampled from 1992-1994 during the dry seasons. Variable length transects laid in different habitats on large islands or across small islands (Devy et al. 1998), where the number and length of transects depended on the size of the island. Information on the numbers of individuals from each transect was not retained, and so we pooled the total numbers of individuals of all species from all transects on a given island for the analyses we present below.

Lizards and frogs were surveyed using bounded quadrats (10 m x 10 m) from November to May 2010–2011 and 2011–2012 (for more details, see Surendran and Vasudevan 2015). Forty-nine quadrats per taxa were placed in rainforests on relatively flat terrain. The number of quadrats sampled was proportional to island size. We used data from 10 islands for frogs, and 11 islands for lizards (we removed islands where either no species were recorded or where only one quadrat was sampled). Here, sample data retained information on the numbers of individuals within each plot, allowing us to calculate patterns of local and regional diversity on each island.

### Hypotheses and analyses

We follow the framework for hypotheses and analyses outlined in Chase et al. 2019 for untangling the potential mechanisms underlying the ISAR for these groups.

First, we estimated the total number of species on each island, which we refer to as S_total_. Because we did not have independent estimates of S_total_ from each island, we combined abundance data from all plots and extrapolated that to an estimated number of species using the Chao1 estimator (Chao 1984, Hseih et al. 2016); this value should be taken as a minimum possible number of species on each island. We then regressed S_total_ against island size to derive an overall ISAR. While useful as a starting point, the relationship between S_total_ and island area cannot be used to go further into dissecting the possible mechanisms underlying the ISAR relationship.

### Can we reject the null hypothesis of random sampling?

We used individual-based rarefaction to evaluate whether the ISAR results deviate from random sampling, or if instead some biological mechanism can be invoked. This approach, similar to the random-placement model of Coleman (1981), uses the individual-based rarefaction curve calculated from all of the transects/quadrats taken from each island. From this island-wide individual-based rarefaction curve, we can then calculate the numbers of species expected for a given number of individuals (n), which we term S_n_. These values (S_n_) were interpolated or extrapolated from the island-wide individual-based rarefaction curves for each island at a common number of individuals (n). In this case, we rarefied S to a reference n, which we calculated as the product of two times the minimum total number of individuals found in an island per dataset (for more details see Chao et al. 2014).

If there is no relationship between S_n_ and island size, then we cannot reject the null hypothesis that the ISAR results from random sampling alone. Alternatively, if S_n_ increases with island size, we can conclude that there is some other mechanism operating that allows more species to co-occur within a given n on larger than smaller islands, which allows us to reject the null hypothesis of random sampling, and indicates that disproportionate effects and/or heterogeneity are playing a role in driving the patterns.

In order to further discern whether any changes in S_n_ were due to changes in the overall evenness of the community, or rather just changes to the rarest species in the community, we calculated a metric of diversity that is primarily sensitive to changes in the most common species, but insensitive to rarer species. Specifically, we used the pooled data to estimate Hurlbert’s (1971) Probability of Interspecific Encounter (PIE),

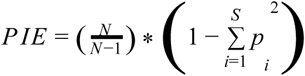

where N is the total number of individuals in the entire community, S is the total number of species in the community, and p_i_ is the proportion of each species i. For analyses, we convert PIE to an effective number of species, S_PIE_ which is described as the number of species that would be observed in a community if all of the species in it were equally abundant (Jost 2006) (S_PIE_= 1/1-PIE, and is proportional to Simpson’s index; Hill 1973, Jost 2006). A relationship between S_PIE_ and island area indicates that larger islands have overall more even abundance distributions. Alternatively, if S_n_ increases with island area, but S_PIE_ does not vary with island area, we would then conclude that only the rarer species are influenced by island area (Chase et al. 2019).

### Within-island β-diversity

A significant relationship between island area and both S_n_ and S_PIE_ can allow us to reject the null hypothesis of random sampling driving the ISAR, but when these values are calculated from pooled data across each island, we cannot differentiate between disproportionate effects and heterogeneity. To disentangle the potential influence of heterogeneity, it is necessary to compare differences in species composition within islands (i.e, β-diversity) that differ in size (Chase et al. 2019). While we only had island-level information on relative abundances for the birds and butterflies, we were able to calculate β-diversity measures from the frog and lizard data where spatially-explicit plot level data were available. To do so, we compared the values of S_n_ when calculated within a single quadrat with the value of S_n_ when calculated from the pooled individuals across all plots. The difference between these two values indicates the degree to which species are clumped in the landscape (i.e., β-diversity). The same can also be done for S_PIE_ to determine whether the clumping is due to more common or rare species. If there is no relationship between either of these β-diversity and island size, we can reject the heterogeneity hypothesis, whereas if measures of β-diversity increases with island size, we can conclude that heterogeneity plays a role underlying the ISAR.

### Statistical analysis

We calculated total estimated species richness (S_total_), the rarefied number of species expected at a common number of individuals (S_n_) and the effective number of species (S_PIE_) using the R package mobr (McGlinn et al. 2019); for lizards and frogs, we calculated these from the pooled data across each island, as well as the plot-level data in order to derive *β* -indices. Code specifically for ISAR analyses are available on GitHub https://github.com/LeanaGooriah/ISAR_analysis. For each taxa, we used linear regression to evaluate the relationship between the various diversity indices (S_total_, S_n_, S_PIE_) and island size.

## Results

Figure 2 illustrates the ISAR relationship for each taxa for each diversity measure and Table 1 gives the regression coefficients. For all four taxa, total species richness (S_total_) increased with island size. Likewise, rarefied species richness, S_n_, increased with island size, allowing us to reject the null hypothesis of random sampling for each taxa. However, we only found a significant increase of S_PIE_, which emphasizes changes in the overall evenness of the community, with island size for butterflies, but not the other three taxa.

**Table 1.** Regression models and their estimates of intercept, slope and R^2^.

| Taxa | Response | $\lg C \pm SE$ | $z \pm SE$ | $R^2$ | $p$-value |
| --- | --- | --- | --- | --- | --- |
| Birds | $\log S_{\text{total}} \sim \log \text{Area}$ | $3.23 \pm 0.08$ | $0.14 \pm 0.03$ | **0.38** | **2.42e-05<sup>9</sup>** |
| | $\log S_n \sim \log \text{Area}$ | $3.07 \pm 0.06$ | $0.08 \pm 0.02$ | **0.31** | **0.0001** |
| | $\log S_{\text{PIE}} \sim \log \text{Area}$ | $2.63 \pm 0.10$ | $0.05 \pm 0.03$ | 0.03 | 0.15 |
| Butterflies | $\log S_{\text{total}} \sim \log \text{Area}$ | $3.04 \pm 0.27$ | $0.14 \pm 0.06$ | **0.39** | **0.05** |
| | $\log S_n \sim \log \text{Area}$ | $2.76 \pm 0.23$ | $0.11 \pm 0.05$ | **0.29** | **0.09** |
| | $\log S_{\text{PIE}} \sim \log \text{Area}$ | $1.82 \pm 0.24$ | $0.17 \pm 0.05$ | **0.53** | **0.02** |
| Lizards | $\log S_{\text{total}} \sim \log \text{Area}$ | $0.64 \pm 0.16$ | $0.21 \pm 0.03$ | **0.77** | **0.0002** |
| | $\log S_n \sim \log \text{Area}$ | $0.75 \pm 0.15$ | $0.16 \pm 0.03$ | **0.67** | **0.001** |
| | $\log S_{\text{PIE}} \sim \log \text{Area}$ | $0.70 \pm 0.18$ | $0.07 \pm 0.04$ | 0.18 | 0.10 |
| Frogs | $\log S_{\text{total}} \sim \log \text{Area}$ | $-0.10 \pm 0.41$ | $0.22 \pm 0.08$ | **0.37** | **0.03** |
| | $\log S_n \sim \log \text{Area}$ | $-0.002 \pm 0.34$ | $0.16 \pm 0.07$ | **0.31** | **0.05** |
| | $\log S_{\text{PIE}} \sim \log \text{Area}$ | $0.44 \pm 0.45$ | $0.046 \pm 0.08$ | -0.13 | 0.62 |

**Figure 1:**
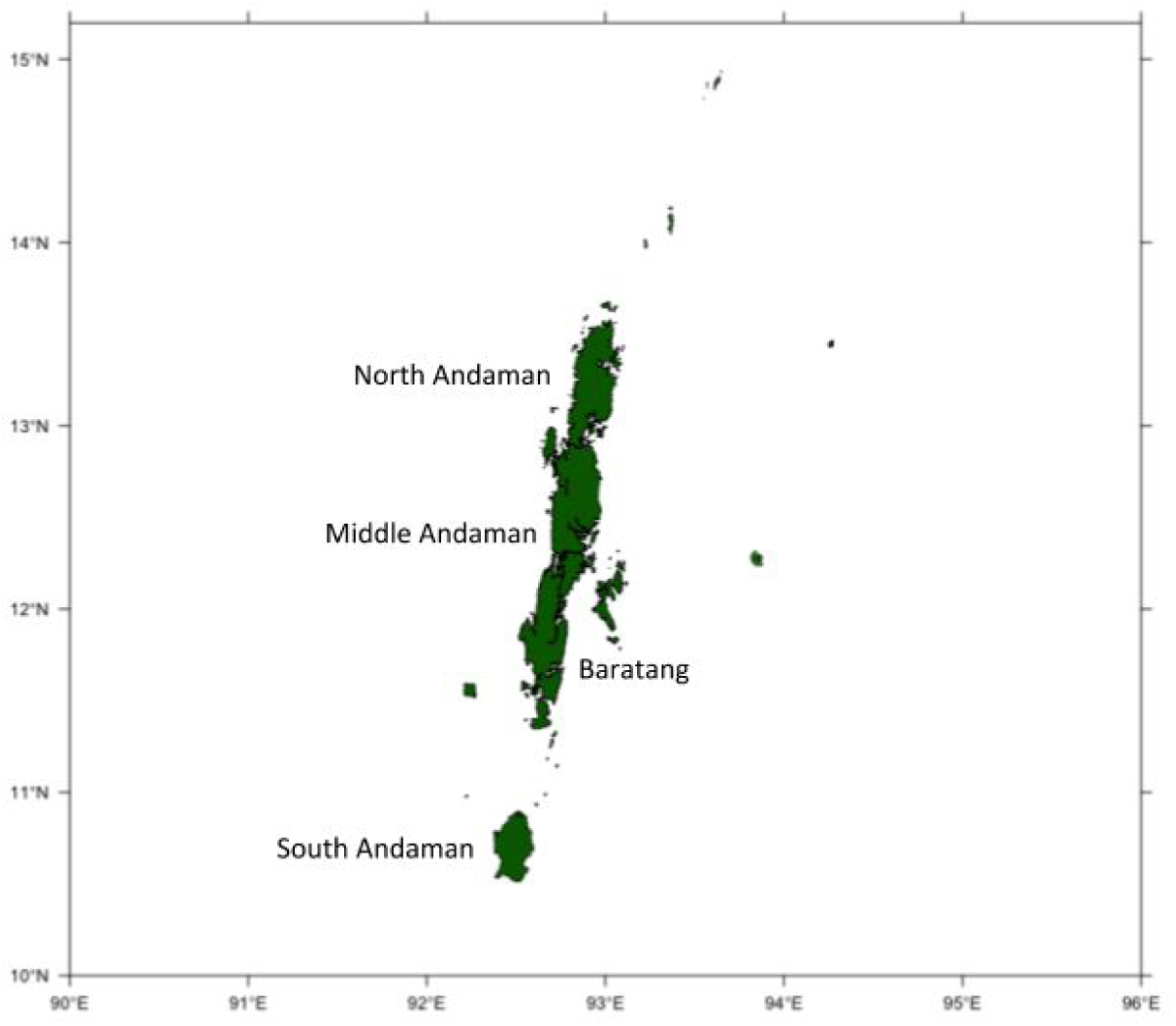
Map of Andaman island group, the four main regions being North, Middle, Baratang and South Andamans.

**Figure 2:**
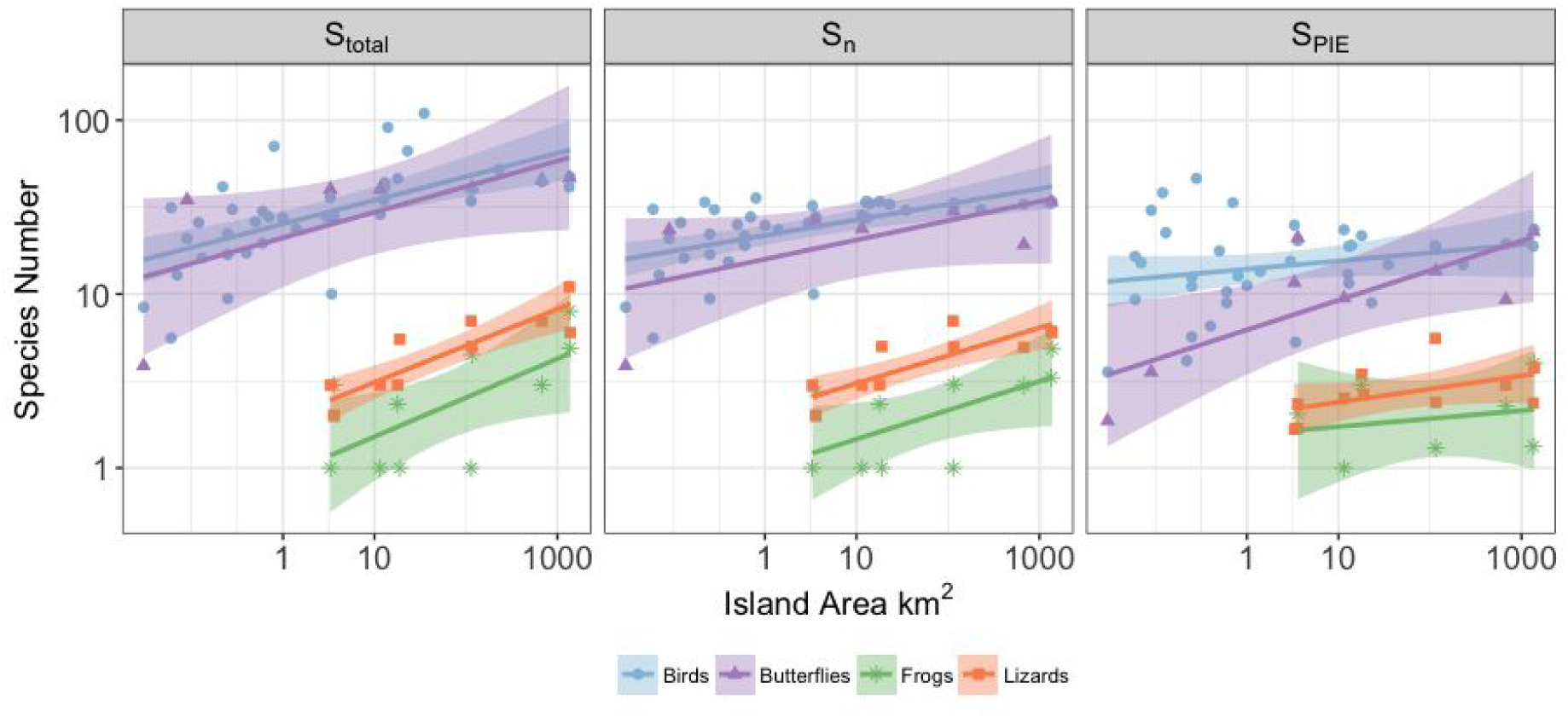
Linear regressions of log-transformed biodiversity metrics against the log of island area (km^2^) for all four taxa. Variables include the total number of species estimated per island from the pooled abundance data (S_total_), the number of species expected at a specific number of individuals (S_n_) and the corresponding effective number of species of the probability of interspecific encounter (S_PIE_).

### Lizards and frogs

For frogs and lizards, we regressed S_n_ and S_PIE_ measured from individual plots (rather than the whole island, as above) against the log of island area and found similar results to the whole-island scale. As a result, we found no difference in either of the β-diversity measures (estimated by taking the regional level estimate divided by the plot-level estimate) with increasing island size for these two taxa (Figure 3, for all four linear regression lines : p-values > 0.1).

**Figure 3:**
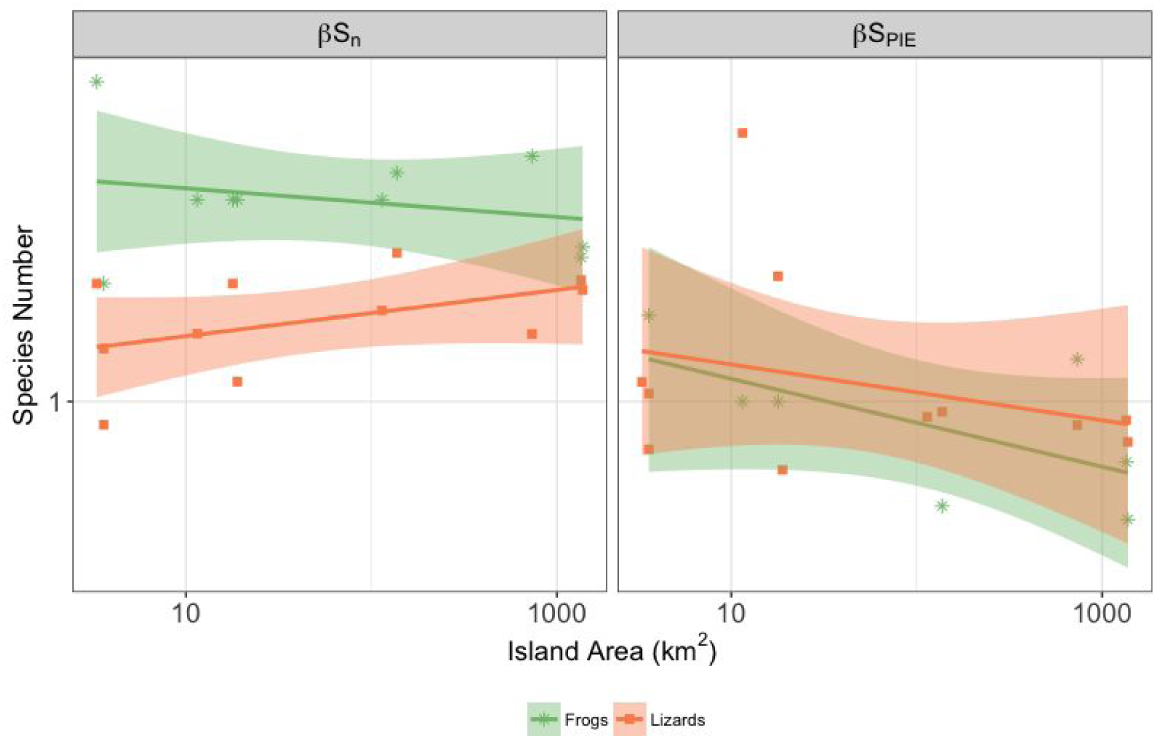
Linear regressions of log-transformed variables S_n_ and S_PIE_ at the *β* -scale against the log of island area (km^2^) for frogs and lizards.

## Discussion

Our results showed that island size had a positive significant effect on bird, butterfly, frog and lizard species richness at the whole island scale (S_total_). This result is not surprising and such a positive ISAR is expected as a result of a number of theoretical expectations and is the most frequently observed pattern (e.g., MacArthur and Wilson 1967, Connor and McCoy 1979, Triantis et al. 2012). However, while significant attention has been paid towards describing the shape of this relationship at the whole island scale, it tells us little about the potential underlying mechanisms of the ISAR.

When we dissected the ISARs of these taxa to discern possible underlying mechanisms, we found an overall consistent pattern that the island-wide rarefied species richness (S_n_) increased with island size. This means more species persist for a given number of individuals than would be expected from a random sampling effect, thus inferring that processes beyond sampling are operating. We used our measure of evenness (the Probability of Interspecific Encounter, PIE), which is relatively insensitive to rare species, and its conversion to an effective number of species to discern whether any changes in Sn were primarily due to an increased probability of rare species persisting beyond sampling expectations on larger islands (in which case, S_PIE_ would not be expected to change), or instead due to changes in both rarer and more common species (in which case, S_PIE_ would increase with island size). For three taxa (birds, frogs and lizards), S_n_ increased with island size, while S_PIE_ did not. From this, we can infer that it was primarily the rarer species that were able to disproportionately persist on larger rather than smaller islands. This could have emerged, for example, because populations on larger islands were more likely to persist by avoiding Allee-effects and/or demographic stochasticity (Hanski and Gyllenberg 1993, Orrock and Watling 2010), or through the increased likelihood of specialized habitats on larger islands (Williams 1964, Kohn and Walsh 1994, Davidar et al. 2001). For butterflies, both S_n_ and S_PIE_ increased with island size, suggesting that not only were rarer species disproportionately favored on larger islands, but that entire shape of the relative abundance distribution became more even on larger islands. Without further information, we cannot explicitly test why butterflies might have differed in their responses to island size compared to the other taxa, but might speculate that owing to their larger population sizes and higher levels of specialization (especially in the larval stage), they were able to more readily alter their relative abundance distributions on larger islands.

Because plot-level data were available for the frogs and lizards, we were able to compare the different biodiversity metrics across scales to explicitly test whether habitat heterogeneity, which would leave a signature in the derived *β* -diversity measures, played a role in driving the ISARs of these taxa. Perhaps surprisingly given the fact that larger islands in this archipelago do have more heterogeneity in habitat types and have a higher proportion of wet evergreen forests that support rarer species (Davidar et al. 2001, Yoganand and Davidar 2000), we found no influence of island size on *β* -diversity of these two taxa despite the fact that they are relatively poor dispersers (Quinn and Harrison 1988, Cook and Quinn 1995, Watling and Donnelly 2006). Thus, at least for these taxa, we can conclude that some mechanism is allowing rarer species to have a higher probability of persistence on larger islands, rather than a mechanism associated with habitat heterogeneity and/or dispersal limitation.

While our results point to a strong influence of island size on both the total number of species (S_total_) as well as the numbers of species persisting when the numbers of individuals are controlled with rarefaction (S_n_), we cannot exclude other variables influencing the species diversity relationships other than area. For example, in a study involving plants on small islands, Panitsa et al. (2006) found strong island species-area relationships but factors such as elevation and the presence of grazing species also explained some of the variance. Another important variable influencing island species-area relationships is isolation, that is, the distance of islands with regard to each other and the mainland (MacArthur and Wilson 1967, Kreft et al 2007). Most of the islands included in our analysis and the Andaman island group in general are quite close to the mainland, so isolation may not have been a likely contributing factor in this case.

In conclusion, we found positive ISARs for all four taxa, but no evidence for sampling effects. These findings suggest that larger islands are important sources of biodiversity, where more species are able to persist than expected from random sampling. Rare species seem to be important drivers of the ISAR, suggesting that rare species are more likely to persist on larger islands either due to disproportionate effects or the availability of more diverse habitats. Moreover, comparing species composition within islands (i.e., β-diversity) can give us additional insight on what drives diversity patterns by allowing us to test for disproportionate vs heterogeneity effects. Overall, our results highlight the importance of larger islands as sources of rare species. This is especially important in nature conservation and planning since smaller islands are usually given higher priority mainly when establishing nature reserves. The protection and presence of nature reserves on larger islands could therefore be a more effective way of protecting rare species from extinction.

## Acknowledgments

We especially thank Jean-Marc Thiollay for supplying the unpublished data. Field work was carried out by K. Yoganand, Jean-Marc Thiollay, M. Soubadra Devy and T. Ganesh with a grant from the Ministry of Environment, France. This work was made possible by the support of the German Centre for Integrative Biodiversity Research (iDiv) Halle-Jena-Leipzig funded by the German Research Foundation (FZT 118).

